# Mosaic of somatic mutations in one of Earth’s largest organisms, Pando

**DOI:** 10.1101/2024.10.19.619233

**Authors:** Rozenn M. Pineau, Karen E. Mock, Jesse Morris, Vachel Kraklow, Andrea Brunelle, Aurore Pageot, William C. Ratcliff, Zachariah Gompert

## Abstract

While evolutionary biology traditionally focuses on the spread of mutations within populations, the dynamics of mutational spread within individuals, particularly in long-lived clonally-spreading organisms, remain poorly understood. Here we examine the genetic structure of ‘Pando’, Earth’s largest known quaking aspen (*Populus tremuloides*) clone. We sequenced over 500 samples across Pando and neighboring clones, including multiple tissue types. At fine spatial scales, we detected significant genetic structure, particularly in leaf tissue, but this signal weakened across larger distances, suggesting either rapid root growth homogenizes the system over time or mechanisms exist that prevent widespread mutation transmission. Phylogenetic analyses date Pando between ∼12,000 and 37,000 years old, supported by continuous aspen pollen presence in nearby lake sediments. Tissues accumulated mutations at different rates, with leaves showing significantly higher mutation loads than roots or branches. This work provides the first quantitative age estimate for this remarkable organism and offers initial insights into the spatial dynamics of somatic mutation in a massive clonal plant. While our reduced-representation sequencing approach limits detection of rare variants, these findings establish a foundation for understanding how long-lived modular organisms accumulate and distribute genetic variation, questions that will benefit from future high-coverage whole-genome sequencing across tissues.

## Introduction

Understanding how mutations arise and spread through a population is essential to understanding biological evolution. The advent of high-throughput genome sequencing has allowed us to study mutational dynamics in a vast array of previously intractable non-model organisms [1], but most prior work has focused on how mutations spread among largely well-individuated organisms (i.e., a life cycle that includes regular genetic bottlenecks), ignoring the effects of within-organism somatic mutations. This is a reasonable assumption for animals, in which germ cells segregate early during ontogeny, but many multicellular organisms (i.e., plants [2–4], fungi [5], red algae [6], brown algae [7]) grow clonally, and do not have germline sequestration.

In contrast to organisms with discrete generations, some organisms grow via clonal extension of a simple modular body plan. This is common within plants and fungi, and can result in the colonization of some habitats over vast distances and exceptionally long periods. For example, seagrasses grow via rhizomal extension, and can have genets that span large areas (e.g., a 47 km transect) [8]. Similarly, a 2,500-year-old clone of the fungus *Armillaria gallica* has spread over 75 hectares of forest rhizosphere [9]. Clonal proliferation may be especially resilient in the face of periodic disturbance. For instance in the Quaking Aspen, *P. tremuloides*, the growth of new ramets is stimulated by nutrients and light availability in areas recently damaged by fire [10, 11]. It is thus not particularly surprising that clonally-expanding organisms can be both very large and long-lived.

Clonally-expanding organisms may evolve quite differently than canonical multicellular organisms that undergo regular unicellular genetic bottlenecks. In the absence of regular bottlenecks, somatic mutations can accumulate and be carried to new tissue as these organisms grow [12]. While the emergence of somatic mutations in animals can lead to lethal cellular proliferation, clonal organisms might be less susceptible to the morbidity and mortality of cancer. Indeed, in organisms like plants, which have permanent cell-cell bonds and a modular growth plan, cancer poses little systemic risk. Notably, clonally-spreading plants and fungi have some of the longest documented lifespans. For example, in seagrasses, such as *Posidonea australis* [13], *P. oceanica* [14], *Thalassia testudinum* [8], or *Zostera marina L*. [15], estimates suggest that individual clones may be more than 6000 years old. With indeterminate growth, the longevity of modules is decoupled from that of the genet (and, in cases where genets remain connected, the organism), making clonally-expanding organisms potentially immortal. In such organisms, novel mutations may even provide a benefit, facilitating adaptive evolution via among-module competition. While some work has examined the dynamics of within-individual somatic mutations [8, 10, 16–18], we still know little about how long-lived clonally-spreading organisms evolve.

Here, we examine the genetic demographic history of Pando, one of the largest genets of the clonally-spreading Quaking Aspen (*P. tremuloides*) [19]. This species can reproduce vegetatively by growing roots, from which new ramets grow. While individual ramet lifespan averages 110 years [20], clones can regenerate from the root stock such that the organism can be far older than its parts. This clone has gathered particular attention for its size (42.6 hectares comprising ∼47,000 individual stems [21]), and is generally thought to be ancient [22]. To investigate its evolutionary history, we sequenced over 500 samples from across Pando and neighboring clones, including leaves, roots, and bark tissues, analyzing patterns of somatic mutations to understand how genetic variation accumulates and spreads within this massive organism. Our phylogenetic analyses, corroborated by local pollen records, date Pando to between 12,000 and 37,000 years old, offering key insights into the evolutionary dynamics of long-lived clonal organisms while providing the first quantitative age estimate for this remarkable organism.

## Materials and methods

### Sampling

To describe the genetic structure and evolutionary history of the Pando clone, we generated four sets of data using different spatial scales and sequencing strategies (Table1). The Pando clone (*Populus tremuloides*) is located in the Fishlake National Forest, Utah, USA (38^*°*^31’N, 111^*°*^45’W), and ranges in altitude from 2,700–2,790 m. To verify the bounds of the Pando clone that had previously been estimated with 7 microsatellite markers [21], we generated a large-scale dataset by sampling leaves from the whole Pando stand, including the neighboring non-Pando clones, on a 50-m grid (sampling done in June 2006 and November 2007, see [21] for more details) (“large-scale dataset”, 184 samples, Supplementary figure S1, left panel). To test for fine-scale within-clone genetic structure, we sampled leaves, roots, bark from the trunk and branches from two additional subsections from within the Pando clone in June 2022 (“fine-scale dataset”, 101 samples, Supplementary figure S1, right panel). One of the sampling sites chosen for this additional sampling is in an area that was clear-cut 30 years ago and the other one is in an older area (Supplementary figures S1 and S2). To avoid batch effects and possible confounding effects of the two different spatial scales, the large and fine-scale datasets were analyzed separately (see ordination plots in Supplementary figure S3). To distinguish somatic mutations in Pando from overall genetic variation in the species, 100 additional leaf samples were collected from *P. tremuloides* in the USA’s Intermountain region (Colorado, Wyoming, Nevada, Idaho) to generate the ‘panel of normals’ (see “Identifying somatic mutations” section). Leaves were kept in paper coin envelope and placed in desiccant. Root and bark samples were placed in polyethylene bags and kept at cool temperatures before long-term storage at -20^*°*^C. Finally, to quantify our ability to accurately identify somatic mutations, we re-sequenced 12 samples from the fine-scale dataset 8 times (same DNA extraction sequenced 8 times, “replicate dataset”, 96 total individual reactions, 80 kept after filtering).

### Sequencing

The 296 leaf samples from the Pando and surrounding clones, and the 45 root samples, 45 leaves and 27 bark samples from trunk and branches were prepared for Genotyping-By-Sequencing (GBS). Woody tissues were powdered using a pestle and mortar and further lysed using TissueLyser II (TissueLyser II, Qiagen). Genomic DNA was extracted using the DNeasy Plant Pro Kit (Cat. No. 69204, Qiagen). To generate a reduced complexity DNA library, the genome was digested using MseI and EcoR1 enzymes. The fragments were labelled and prepared for sequencing using oligonucleotides consisting of Illumina adaptors and unique 8-10 base pair (bp) sequences. The fragments were amplified and size-selected to keep fragments between 300 and 400 base pairs. Library preparation and sequencing were done in three batches, with 367 samples sequenced with an Illumina HiSeq 4000 (1 × 100 base pair reads) in 2018, 126 and 96 samples sequenced on a NovaSeq (1 × 100 base pair reads) in 2022 and 2024, respectively (one lane each). The 2018 and 2022 samples were sequenced at the University of Texas Genomic Sequencing and Analysis Facility (Austin, TX, USA). The 2024 samples were sequenced at the Utah State Genomics Core facility. The total number of reads was 1,027,955,624. The mean sequencing depth was 13.5× for the large-scale dataset, 18.1× for the fine-scale dataset and 43.4× for the replicates.

### Genome alignment and variant calling

The raw fastq files were filtered by removing PhiX and adapter sequences. We matched each barcode to their corresponding sample ID and split the reads to their individual sample files. We used the mem algorithm from bwa (default options, version 0.7.17-r1188, [23]) to align the reads to the published reference genome for *P. tremuloides* [24], and samtools to compress, sort and index the alignments (Version:1.16 [23]). We called the variants using samtools mpileup algorithm (Version: 1.16) and bcftools (Version: 1.16). The large-scale and fine-scale datasets were pooled for variant calling, and the replicate and “panel of normals” datasets were kept separate. We kept mapped reads with a quality *>*30, skipped bases with base quality *>*30 and ignored insertion–deletion polymorphisms. At this step, we separated the fine-scale and large-scale samples. We applied a set of filters to all variant sets (three datasets with both somatic and germline mutations) by keeping the sites for which we had data (mapped reads) in no fewer than 60% of individuals, a mean coverage per sample of 4× minimum, and at least one read supporting the non-reference allele. We also removed SNPs failing the base quality rank-sum test (*P <* 0.005), mapping-quality rank-sum test (*P <* 0.005), and the read position rank-sum test (*P <* 0.01).

### Delineating the Pando clone

In order to differentiate between the samples pertaining to the Pando clone and the surrounding clones in the large-scale dataset, we obtained Bayesian estimates of genotypes for the full set of SNPs identified above (somatic and germline mutations). We specifically computed the posterior mean genotype as a point estimate based on the genotype likelihood from bcftools and a binomial prior based on the allele frequency estimates from the vcf file (this posterior has an analytical solution). We used principal component analysis (PCA) to ordinate the samples; this was performed on the matrix of centered but not scaled genotype estimates. The PCA separated the Pando clone samples, from the surrounding clone samples (Figure 1) in PCA space. We used k-means clustering (R kmeans function, with K=2) to identify and label the different clusters of samples (Pando and non-Pando samples). There were 9,424 variants in the Pando variant file at this stage.

**Fig 1.**
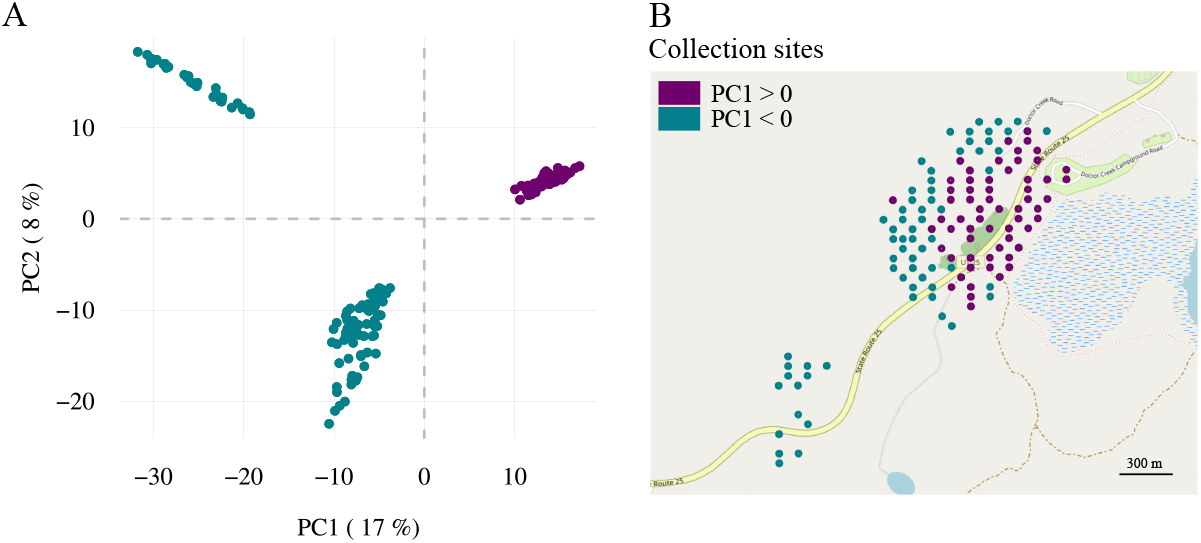
Parsing out the Pando samples from the surrounding clone samples. (A) The projection of genotypes (22,888 variants) form three distinct clusters: two clusters with negative PC1 values and one cluster with positive PC1 values. Points are labeled with a color proportional to their PC1 value. (B) Plotting the PC1 value into the sampling space delineates the Pando cluster (positive PC1 values) from the surrounding clone clusters (negative PC1 values).

### Identifying Somatic Mutations

Germline mutations are inherited and should be common to Pando as a whole. Somatic mutations, however, are mutations that appeared after seed formation and during the organism’s growth, potentially capturing the evolutionary history of the organism. To identify somatic mutations in the Pando clone, we developed a comparative analysis pipeline using neighboring clones and regional samples as reference points. We created a “panel of normals” [25] reference dataset comprising variants from neighboring clones and 100 *P. tremuloides* samples collected across the USA’s Intermountain region (Colorado, Wyoming, Nevada, Idaho). We implemented multiple filtering steps to ensure accurate mutation detection. First, we removed variants present in both the Pando samples and the “panel of normals” reference dataset, as these likely represented germline mutations or highly mutable sites. Second, to minimize false positives from sequencing errors given the inherent per base pair error rate of approximately 0.31% for Illumina reads [26], we excluded mutations detected in only one sample. Third, to validate our mutation detection approach, we performed technical replication by sequencing 12 samples eight times each from the same DNA extraction. In the replicate dataset, we classified mutations as somatic if they appeared in at least two replicates per sample but in no more than 80% of total samples, with this threshold chosen to exclude potential germline mutations that might not be detected in all samples due to technical limitations. Finally, in the fine-scale and large-scale datasets, we classified mutations as somatic if they appeared in at least two samples, but in no more than 80% of total samples. For a full description of our mutation filtering pipeline, see Supplementary Methods.

### Benefits and limitations of reduced-representation sequencing

Reduced-representation sequencing like GBS enabled high sample throughput and broad spatial coverage, but its reduced genomic representation and moderate depth limit sensitivity to rare somatic mutations in a triploid genome. The average sequencing depth across samples was ∼13-14×, with a per-site minimum threshold of 4×. While sufficient for genome-wide patterns, this depth is suboptimal for confident detection of low variant-allele-frequency mutations in triploids. Additional uncertainty arises from GBS–specific errors and limitations of the reference genome, which can increase mis-mapping and obscure true signal. We explicitly quantified error rates and applied conservative variant calling, likely underestimating the number of somatic mutations. Higher-coverage whole-genome sequencing, and ideally single-cell sequencing of defined meristem lineages, will be needed to refine mutational and evolutionary parameter estimates in this iconic organism.

### Spatial analyses

To detect spatial structure in the dataset, we applied the same set of analyses on two different datasets: (1) a large-scale, and (2) a fine-scale dataset. We first compared the proportion of shared variants per pair of samples to their physical distance (number of shared mutations between a pair of samples, divided by the mean number of mutations for the same pair of samples). We then computed the mean geographic distance between groups of samples sharing a mutation (with the mean taken over all mutations). We used Vincenty ellipsoid method (distVincentyEllipsoid function in R) to calculate the shortest spatial distance between two samples. For each analysis, we compared the empirical values to values obtained from a randomized dataset to assess the significance of the results. To generate null distributions, we randomized either the genotypes or the pair of spatial coordinates (latitude and longitude) and ran the same analysis as ran on the non-permuted data (500 or 1,000 permutations).

### Coalescent model using BEAST

We used the software package BEAST (version 2.7.5) to estimate the height of the phylogenetic tree for the Pando samples based on the accumulated somatic mutations; this was done using a coalescent Bayesian skyline model for effective population size [27–29]. We chose the GTR nucleotide-substitution model to account for unequal substitutions rates between bases [30], starting with equal rates for all substitutions. We selected an optimized relaxed clock with a mean clock rate of 1. To generate the nexus file, we assigned genotypes based on the point estimate value (the sample is heterozygote, coded “A”, for the mutation if the point estimate value *>* 0.5, homozygote otherwise, coded “T”). A single chain was run for 7 × 10^7^ states. To estimate the age of the tree, we converted the phylogeny height to years *a posteriori* following this calculation:

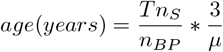

with *T* being the phylogenetic tree height as given by BEAST, *n*_*S*_ the total number of mutations, *n*_*BP*_, the total number of base pairs sequenced, *µ* the leaf somatic mutation rate (1.33 ∗10^−10^ per base per haploid genome per year [31]), taking into account that the Pando clone is triploid [32, 33]. The total number of base pairs sequenced (129,194,577) was estimated using angsd [34], and reduced following the proportion of base pairs that we filtered out because of low coverage (48%).

### Accounting for missing mutations

To estimate how many somatic mutations we may have missed, we combined the 12 samples from the fine scale dataset, with the replicate dataset (these same 12 samples sequenced 8 times) and performed joint variant calling, identifying 55 somatic mutations. Comparing detection in the replicate set versus the original single-pass dataset showed that, on average, mutations present in the replicate set were recovered about 20% of the time in the original dataset. This result was robust to permuting the reference and replicate set samples (Supplementary figure S9). Given this detection rate, our divergence-time estimates explicitly bracket the extremes: either we are missing roughly 80% of true somatic mutations or only 20% of the mutations we detect are true positives. To account for those uncertainties when calculating the age of the Pando clone, we considered three different scenarios : (i) all mutations detected are real, (ii) only 20% of the mutations we detect are real and (iii) 80% of true mutations are missing. To quantify how missing mutations affect phylogeny height, we randomly censored increasing proportions of mutations, inferred trees in BEAST. We leveraged the linear relationship between the fraction of mutations missing and the phylogenetic tree height to estimate the Pando clone age. While this approach accounts for the extremes and stabilizes the divergence-time estimates, distinguishing among these possibilities and narrowing the uncertainty will require extremely high-coverage whole-genome sequencing.

### Pollen analysis

Pollen analysis followed standard acid digestion procedures [35]. Pollen residues were classified and tabulated using light microscopy at 40×until a minimum of 300 terrestrial grains were counted. Pollen identification was assisted by relevant keys and literature (e.g., Kapp et al. 2000 [36]). We assume that the *Populus* pollen type, which is generally not diagnostic to species-level assignment, reflects quaking aspen in this environmental setting.

## Results

### Delineating the Pando clone

Pando samples (89 out of 184 samples) formed a distinct cluster in PCA space (Figure 1A) with spatial boundaries for Pando that were consistent with previously defined clone boundaries based on morphological differences [19], and microsatellite markers [21, 32] (Figure 1B and Supplementary Table 1). We thus verified the spatial extent, 42.6 ha, of Pando.

### Identifying somatic mutations

We generated a replicate dataset to quantify our ability to recover rare mutations between samples. Our filtering approach (see Methods for details) identified a set of 101 putative somatic mutations, each of which was present in less than 40% of samples (no mutations were detected between 40% and our 80% threshold; Figure 2A,B). We found that when a mutation was detected in at least two replicates of a sample, it appeared on average in 3.5 replicates total (44% of replicates), significantly higher than expected by chance (randomization test, null expectation = 0.37 with 1,000 permutations, P *<* 0.001, Figure 2C). The detection of mutations remained consistent across varying coverage depths (Supplementary Figures S4 and S8). While these results validate our ability to detect somatic mutations, they also suggest our approach may be missing some true mutations, a limitation we address in our age estimation analyses. Having established our mutation detection methodology, we applied these filtering criteria to both our large-scale and fine-scale datasets for subsequent analyses.

**Fig 2.**
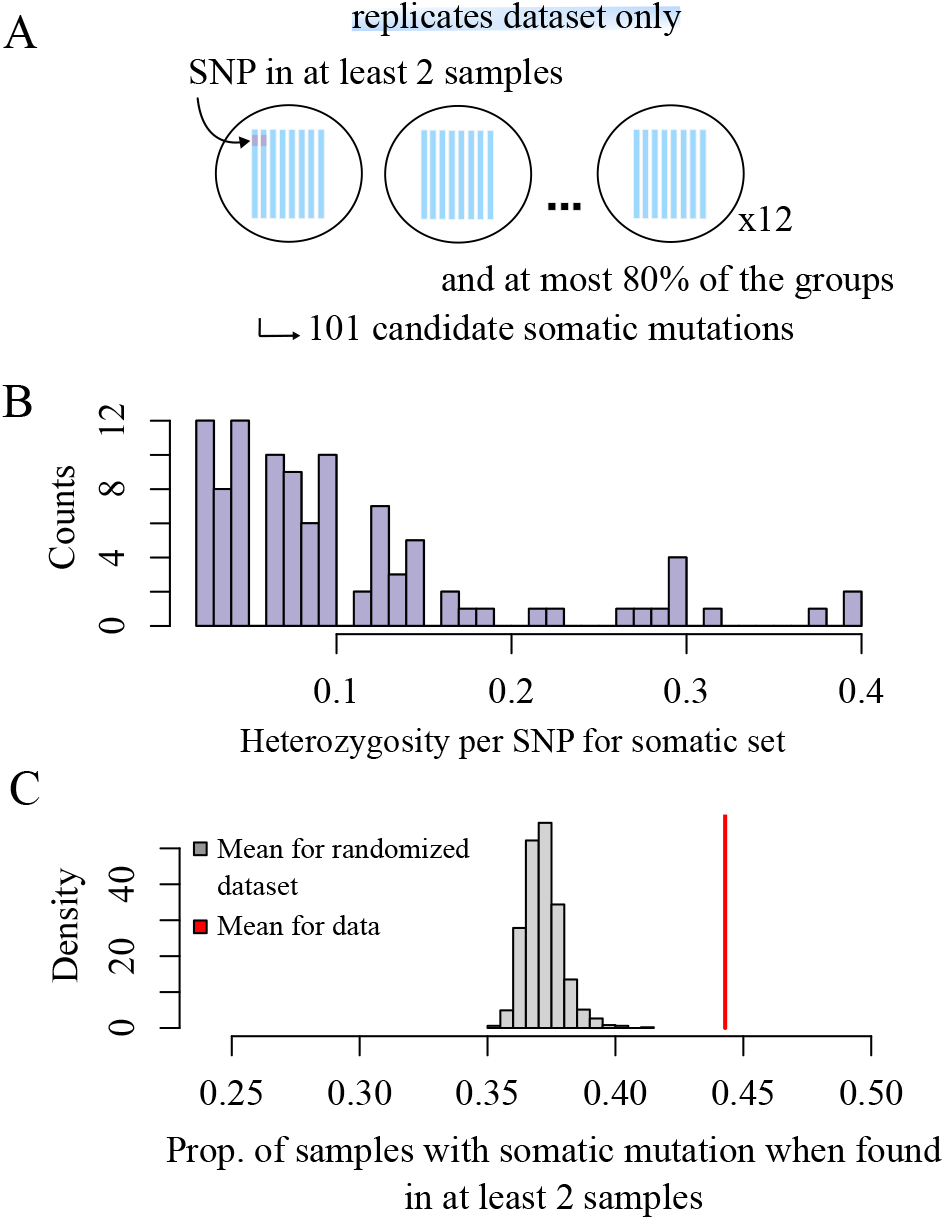
Replication power for somatic mutations. (A) To filter for somatic mutations, we labeled the mutations that were found in at least two samples per replicate group, and at most 80% of the samples (see methods for details on the filters). We identified 101 somatic mutations, (B) found in less than 40% of the individuals. (C) If a mutation is present in two samples in a group, it is found on average in 44% of the samples total.

### Patterns of spatial genetic structure for somatic mutations - large-scale

We identified 3,942 putative somatic mutations from the 89 Pando ramet samples (large-scale dataset, Table 1). On average, samples shared 26.8% somatic mutations (range = 583 to 1,679 mutations, or 14.8–34.7%). Due to clonal reproduction and spatial restriction in dispersal, we expected to observe a non-random spatial distribution of somatic mutations [37]. More specifically, we expected ramets that are close in space to share more mutations than ramets that are further apart from each other. However, there was only a weak correlation between the proportion of shared variants and the physical distance between pairs of ramets (Figure 3A, Pearson correlation coefficient = −0.02, 95%*CI* = [ −0.05, 0.00], Figure 3B, null expectation = -0.001 with 1,000 permutations of the somatic mutation set, *P <* 0.001). We uncovered further spatial structure when focusing on spatial distribution of each somatic mutation. The mean distance between all samples sharing a mutation, averaged over all mutations, is smaller than expected by chance (Figure 3C&D, mean distance for groups sharing a somatic mutations is 264.28 m, as compared to the mean distance (null expectation) of 279.93 m for a randomized dataset with 500 permutations of the sample coordinates, *P <* 0.002). Given that a single root can extend up to 15 m [38], and our grid sampling had a minimum distance of 50 m, we hypothesized that we might be missing spatial signals at finer scales. Additionally, focusing solely on leaves could overlook somatic mutation signals, as clonal aspen expand through their roots (Figure 4). To better understand the spread of somatic mutations within and between ramets and tissue types, we conducted our analyses at a finer spatial scale by comparing samples from sub-sections of the clone and from different tissues within ramets.

**Table 1.** To study the evolutionary history of the Pando clone, we generated datasets at different spatial scales and using different sequencing strategies. The large-scale and fine-scale datasets have the same initial number of mutations as the variant calling was done on both sets at once. Nb. = number.

| Dataset | Nb. of samples | Nb. of variants<br>(all/somatic) |
| --- | --- | --- |
| large-scale | 184 (Pando + neighboring clones)<br>89 (Pando only) | 22,888/-<br>15,925/3,942 |
| fine-scale | 101 | 15,925/3,034 |
| replicate | 80 | 4,607/101 |
| panel of normals | 100 | 99,234 |

**Fig 3.**
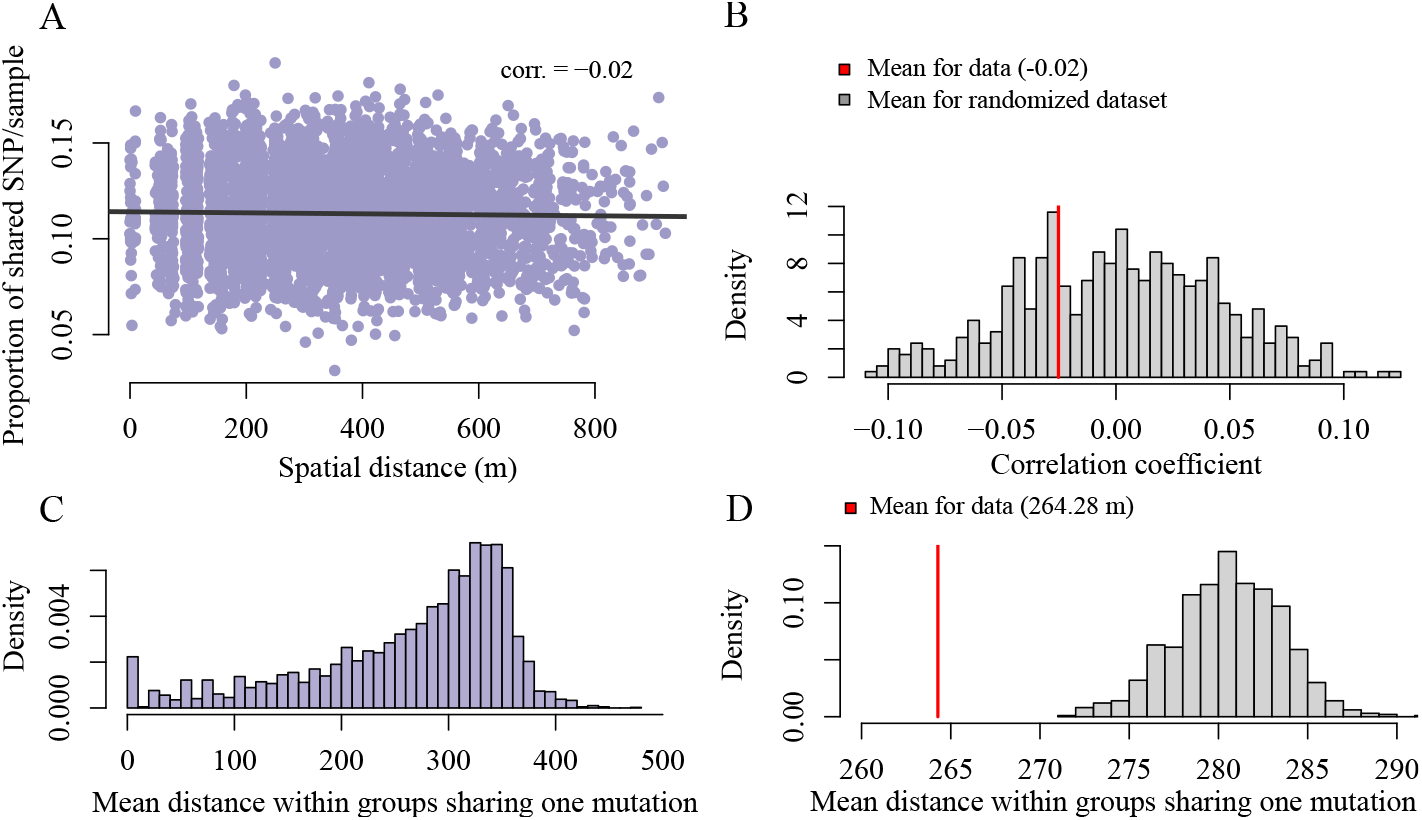
Detecting spatial genetic structure at large scale. (A) We use the set of 3,942 somatic mutations identified in the Pando clone samples to test for spatial genetic structure. Focusing on the sample-level, we observe that the number of shared variants between pairs of samples decreases with the physical distance between sample pairs (Pearson correlation coefficient between number of variants and spatial distance is −0.02, 95% CI = [ −0.05, 0.00]), which is significantly different from a randomized distribution (*P <* 0.001) (B). (C & D) Focusing on the variant-level, we find that the mean distance within a group of samples sharing the variant is significantly less than expected by chance (mean distance for data is 264.28 m and mean distance for randomized dataset is 279.93 m, *P <* 0.001).

**Fig 4.**
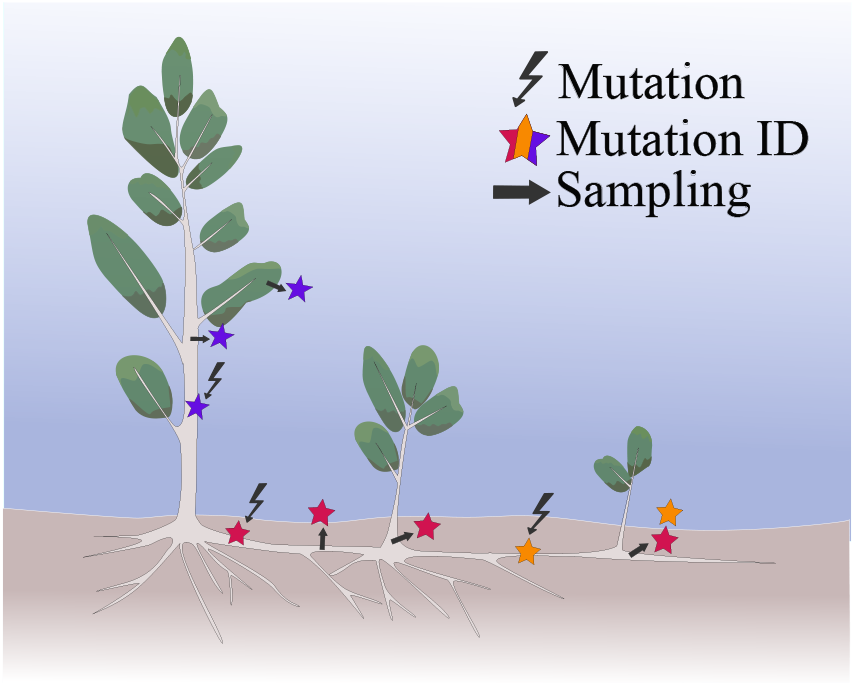
Conceptual model of somatic mutation inheritance between ramets within an aspen clone. When a mutation arises, we expect it to propagate down to the new tissues as the clone continues to grow. New mutations are symbolized with the lightning bolt. The mutation identity is marked as a colored star and the dark marks corresponds to where samples could be collected from the clone.

### Patterns of spatial genetic structure for somatic mutations - fine-scale

Overall, we found significant evidence of genetic structure in our set of 3,034 mutations, with the proportion of shared mutations decreasing with spatial distance (Figure 5A,Pearson correlation coefficient = -0.1, 95% CI = [-0.12, -0.07], null expectation = 0.00 with 500 permutations, *P* = 0.006). The signal was especially strong for leaves (Pearson correlation coefficient −0.44, 95% CI = [−0.49, 0.38]), with more somatic mutations shared between spatially close leaves compared to random (*P <* 0.001). The roots also shared significantly more mutations than expected under a null distribution (Pearson correlation coefficient−0.11, 95% CI = [−0.18, −0.03], *P* = 0.026). This signal was not observed in the branches and the shoots (Pearson correlation coefficient 0.06, 95% CI = [−0.24, 0.11] for branches and −0.05, 95% CI = [ −0.37, 0.28] for shoots).

**Fig 5.**
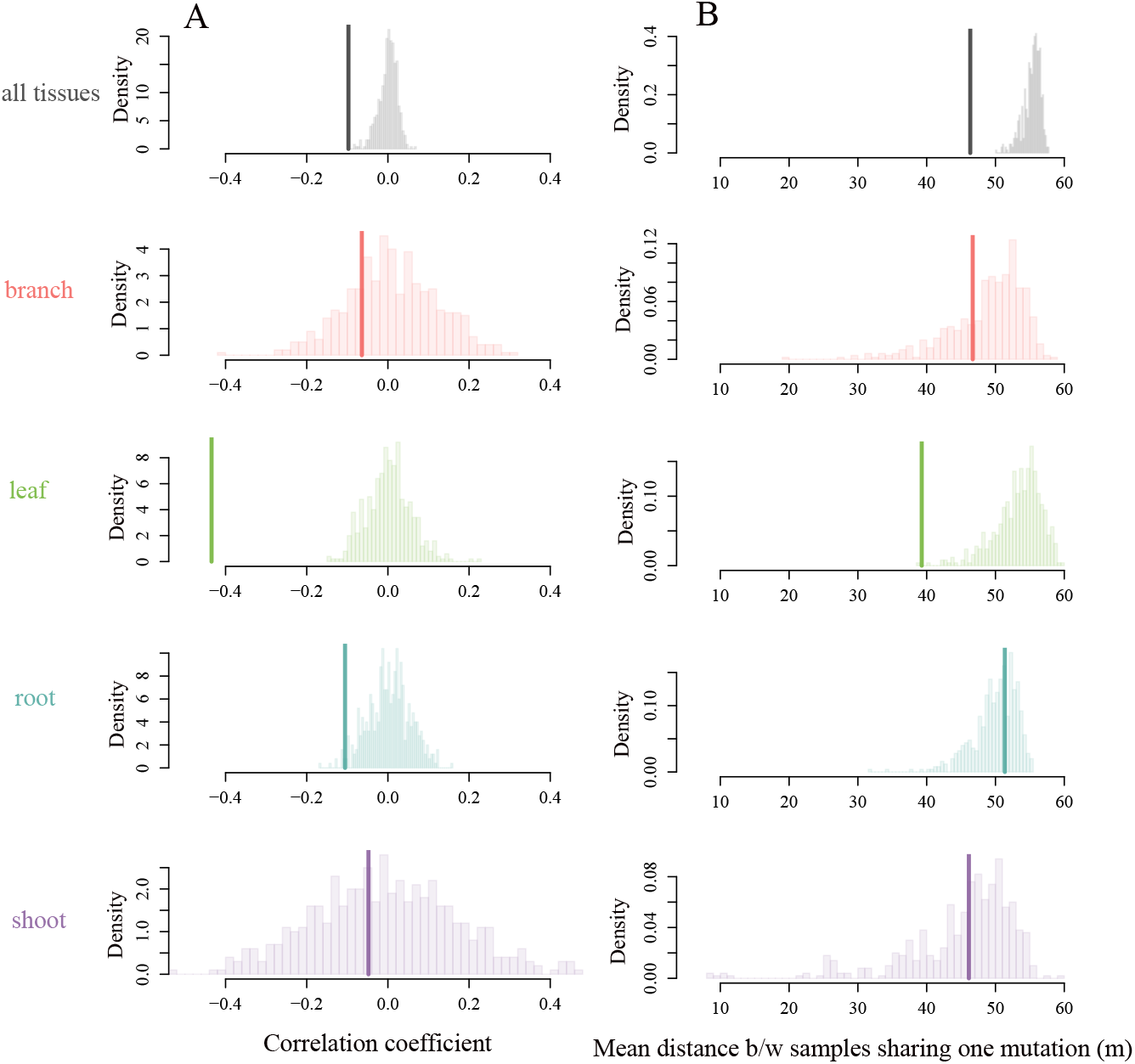
Detecting spatial genetic structure at the finer scale. We use the set of 3034 somatic mutations detected in the finer scale dataset to test for smaller-scale and within tissues spatial genetic structure. (A) Focusing at the sample-level, we observe an overall significantly negative correlation between genetic and physical distance (thick lines, Pearson correlation coefficient = −0.097, [*CI*] = [−0.12, 0.07]), driven mostly by the leaves and the roots (compared to null distributions, *P <* 0.001 and *P* = 0.026, respectively). (B) Focusing on the variant-level, we find that the mean distance within a group of samples sharing the variant (thick line, mean distance for the data is 46.33 m) is significantly less than expected by chance when considering all tissue types together (mean distance for the null distribution is 55.31 m, *P <* 0.001), signal that is mostly driven by the leaves (mean distance for leaves only is 39.28 m, as compared to 53.36 m expected under the null distribution, see S7 for means and p-values).

A variant-level approach showed that the number of shared somatic mutations per pair of samples decreased with increasing spatial distance (Figure 5B, mean distance for groups sharing a somatic mutations is 46.33 m, as compared to the mean distance (null expectation) of 55.31 m for a randomized dataset with 500 permutations, *P* = 0.002). The leaves showed the strongest spatial structure signal using this metric (Figure 5B and Supplementary figure S5), while other tissue types did not differ from the null expectation. The absence of signal in the shoots and branches may be partly explained by the significantly higher number of mutations recovered in leaves compared to other tissues (Supplementary figure S6). Importantly, this spatial structure in the leaves would not be expected if the putative somatic mutations were primarily sequencing errors (which should be randomly distributed with respect to distance) or germline variants (which should be broadly shared across ramets regardless of proximity). Thus, the observed clustering provides validation that we are detecting somatic mosaicism, and strengthens confidence in the biological signal captured by this dataset.

### Age of the Pando clone

We reconstructed the phylogenetic history of the Pando samples with a variable population size coalescent model (BEAST2 [27]). To take into account the effect of missing mutations on the phylogenetic tree height and thus the Pando clone age, we empirically estimated the relationship between the proportion of missing mutations and the phylogenetic tree height (Figure 6A). This relationship was linear, which we extrapolated to take into account false negatives or positives (i.e. mutations that we either missed, or called but are not real). This scaled tree height was then converted to years (see Methods for more details).

**Fig 6.**
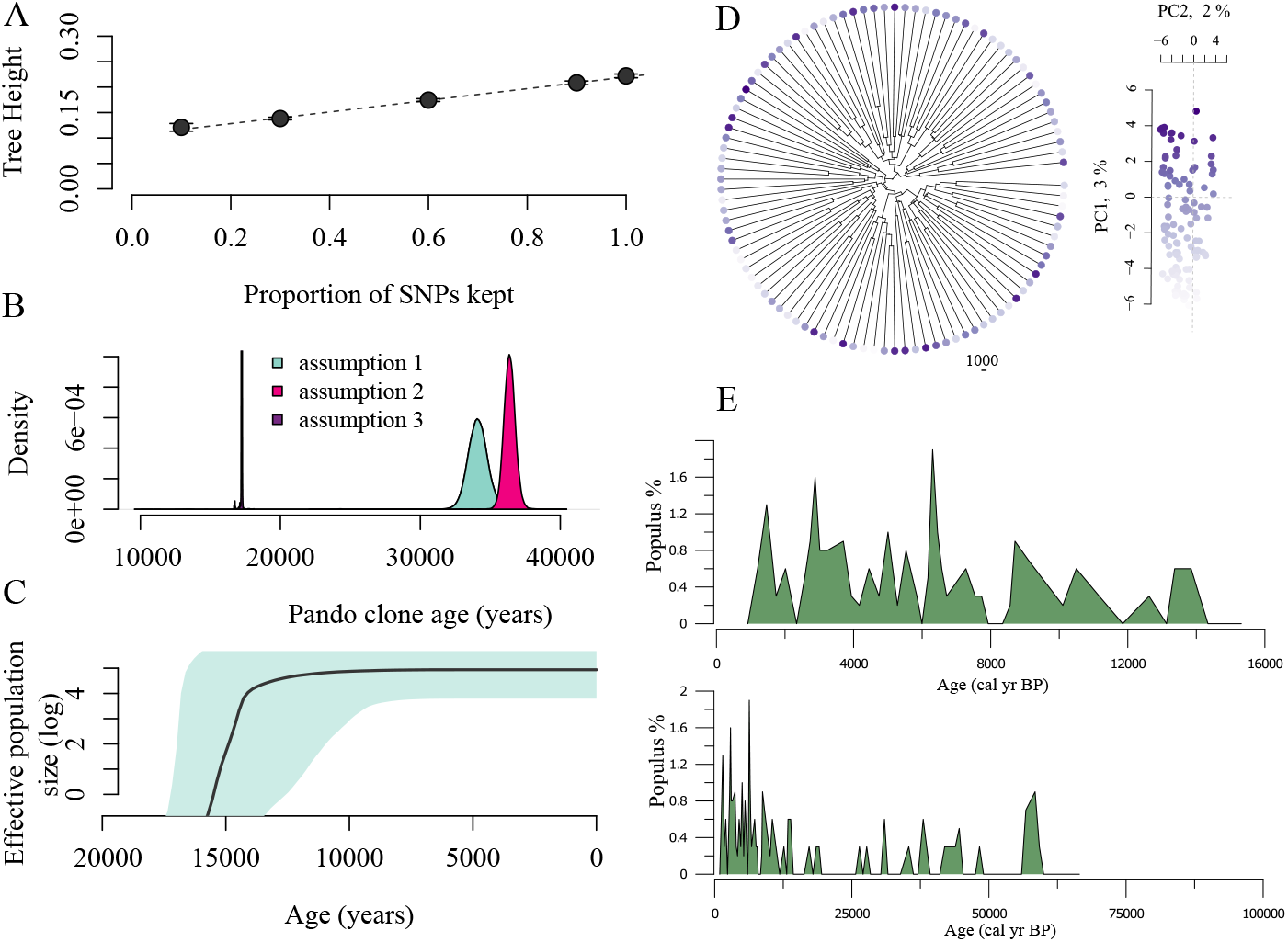
Pando is between 17,000 and 36,000 years old based on the large-scale dataset. (A) We use the relationship between the proportion of missing mutations from a simulated dataset and the phylogenetic tree height to take into account the somatic mutations that we might be missing in the Pando clone (linear regression *y* = 0.10 + 0.11*x, P <* 2.2∗ 10^16^, *R*^2^ = 0.92). (B) With this correction, we calculate the Pando clone age based on three different assumptions: (1) if the mutations we detect are all real, the Pando clone would be about 34,055 years old (±sd = 1,255 years); (2) if we are missing 80% of the mutations, then the clone would on average be 36,351 years old (±sd = 733 years); (3) finally, if only 20% of the mutations we detect are real somatic mutations, the Pando clone would be 17,238 years old (±sd = 29 years). (C) The Bayesian skyline plot suggests a steady population increase followed by a plateau. Note that this example was scaled for assumption 1 (all the mutations that we detect are real somatic mutations). (D) Despite thousands of years of evolutionary history, the Pando clone shows minimal phylogenetic structure (points colored according to PC1 score). (E) Pollen records from the Fish Lake show *Populus* was consistently present during the last 15,000 years, and generally well-represented over the last 60,000 years.

We calculated three different estimates of Pando’s age based on three different assumptions (Figure 6B). First, if the mutations we detected are all true positives and we did not miss any somatic mutations in the proportion of the genome we sequenced, we do not have to apply any correction to the phylogeny height conversion and the Pando clone would be about 34,055 years old (assumption 1, sd = 1,255 years). Second, if we take into account that we only detected 20% of the somatic mutations present in the samples and use the linear relationship (Figure 6A) to account for false negatives, then the clone would on average be 36,351 years old (assumption 2, sd = 733 years).

Finally, if only 20% of the mutations we detect are true positives, the Pando clone would be 17,238 years old (assumption 3, sd = 29 years). The population dynamics reconstruction suggest a slow and steady increase during the first half of Pando’s life, followed by a steadier population size (Figure 6C). The unit of effective population size here can be thought of in terms of cellular lineages giving rise to new tissues (as compared to individuals when working with germline mutations). Despite its thousands of years of history, the phylogeny of the Pando clone samples suggests only minimal structure (Figure 6D). The same analysis of the fine-scale dataset suggests results of a similar range, that is, an age for Pando between ∼12,000 and 37,000 years if we merge the range estimation given from both fine-scale and large-scale datasets (Supplementary figure S10). Interestingly, pollen records from the Fish Lake support a continuous presence of *Populus* during the last 15,000 years, potentially up to 60,000 years ago,which generally coincides with our age estimates for Pando (Figure 6E).

## Discussion

In this study, we explored the evolutionary history of one of Earth’s largest clonal organisms, confirming that the Pando clone of quaking aspen consists of a single genet spanning 42.6 hectares. Using reduced-representation sequencing (GBS), we generated the first large-scale genomic assessment of somatic mutations in this ancient clone. Because GBS samples only a fraction of the genome and at moderate sequencing depth,and because Pando is triploid, our ability to detect rare somatic variants is limited. We quantified uncertainties using technical replicates and sensitivity analyses. We find that somatic variants exhibit fine-scale spatial patterns, especially in leaves and roots; patterns that would not be expected from random sequencing errors or from germline variants—supporting genuine somatic mosaicism. Incorporating potential missing mutations into our phylogenetic analyses, we estimate that Pando is at least 12,000 years old (Figure 6), with upper bounds depending on the assumed rate of undetected mutations. Our divergence-time inferences bracket extreme assumptions about error and missingness and are therefore robust. Reducing uncertainty in timing and fine-scale structure will require high-coverage whole-genome, and ideally single-lineage, sequencing.

Pando could not have persisted if glaciers occupied Fish Lake Plateau. Boulder exposure ages from the plateau suggest a local last glacial maximum of 21,100 years ago, raising the possibility of mountain glacier coverage in the area [39]. However, models of Fish Lake Plateau glaciers show they occurred at elevations between 2,950 and 3,190 m— notably higher than Pando’s current location of 2,700 m [39]. This suggests that vegetation could have persisted at Pando’s location throughout this glacial period [40, 41]. Supporting this interpretation, subfossil pollen analysis from nearby Fish Lake sediment cores demonstrates a continuous presence of *Populus* pollen over the last 15,000 years, with evidence of its presence extending back approximately 60,000 years (Figure 6E, upper panel).

To explore the spatial genetic demography of Pando, we sequenced leaves across a 50-m grid covering the entire Pando area as well as leaves, branches, shoots and roots at a finer scale, with samples collected 1-15 m apart in two locations within the clone. Our findings reveal spatial genetic structure within the clone, with samples sharing more mutations when geographically closer (Figure 3 & 5). While we were able to detect this spatial signal at a fine-scale in the leaves and roots, it was weaker at larger scales than expected and usually observed in clonal organisms [37, 42, 43]. Although we can clearly distinguish Pando samples from neighboring clones (Figure 1) and detect some internal genetic structure within the clone (Figures 3 & 5), the relatively low number of shared mutations between closely related tissues (roots, shoots and branches, Figure 5) suggests an intriguing underlying dynamic.

The weak spatial genetic structure we observed could be explained by two non-mutually exclusive hypotheses. First, given that aspen roots can grow up to 15 meters per generation [38], and Pando spans roughly 2 kilometers at its widest point, it would take only about 133 generations of ramet growth for a lineage to traverse the entire clone. Over thousands of years, this relatively rapid growth potential could lead to Pando functioning as a well-mixed system, effectively homogenizing genetic variation across the organism. Alternatively, research on within-clone mutation diversity suggests a different explanation: somatic mutations may not always be passed on asexually-produced offspring [44]. This pattern has been observed in *Fragaria vesca* (woodland strawberry), where mutations present in mother plants were absent in daughter plants propagated via stolons [16]. Such observations suggest that somatic mutations occurring in local tissues are not always passed down to the next generation of cells. As roots grow, the meristematic island that will give rise to new ramets gets pushed by waves of cells, protecting the stem cells from mutation accumulation [45].

The detection of somatic mutations in plants is further challenged with the fact that they are layer-specific [46]. Despite prolonged lifespan and exposure to significant environmental changes, plants seem to have evolved mechanisms that protect the meristems from accumulating mutations [18]. When sequencing entire tissues, we might be observing the localized accumulation of somatic mutations rather than the cell lineages contributing to organismal evolution, which would explain the relatively weak spatial genetic structure.

Our results suggest differing rates of somatic mutation accumulation between tissues that constitute the “germline” (i.e., contribute to future generations of ramets) versus those that do not, and between annual and perennial tissues. We found that leaves accumulate more mutations than branches, shoots, and roots. This pattern is consistent with findings from other plant systems: in *Fragaria vesca*, for instance, petals (which are even more ephemeral than leaves) showed higher mutation rates than leaves [16]. Similarly, in *Prunus mira* (peach trees), mutation accumulation in branches, which are involved in sexual reproduction, was lower than in roots [16]. The general pattern emerging from these studies is that tissues with higher cell division rates or shorter lifespans tend to accumulate more mutations.

These tissue-specific differences in mutation accumulation have important implications for our age estimates of the Pando clone. Our dating approach necessarily relied on mutation rates derived from leaf tissue to estimate mutation accumulation across the entire organism. However, given that leaves show higher mutation rates compared to other tissues, as demonstrated both in our results and in other organisms where mutation rates can vary by an order of magnitude across tissues [47], our age estimates likely represent conservative lower bounds. The challenge of accurate age estimation is further complicated by cellular mosaicism within tissues. While emerging single-cell sequencing methods [48] may eventually enable more precise tracking of mutation lineages in non-model organisms like aspen, current technical limitations necessitate working with tissue-level data that potentially masks finer-scale genetic variation.

This study provides new insights into the evolutionary history of Pando, one of Earth’s oldest and largest known organisms. By analyzing somatic mutations across different spatial scales and tissue types, we estimate the clone’s age to be at least 12,000 years old, with upper estimates reaching up to 37,000 years. Our findings reveal a weaker-than-expected spatial genetic structure within the clone, suggesting that Pando is either well mixed due to rapid growth, or that mutations accumulate locally rather than dispersing consistently along tissue lineages. The latter would support the hypothesis that mutation-protected pools of cells help maintain the genetic integrity of an indefinitely growing organism. In addition, this work underscores the technical challenges of studying rare mutations in non-model organisms, and paves the way for future studies on mutational dynamics in clonally-spreading long-lived perennials.

## Contributions

RMP, KM, ZG and WCR conceived the study. RMP, AP and KM sampled the Pando clone data. JM, VK and AB sampled and analyzed the pollen data. RMP and ZG performed the analyses. RMP, ZG and WCR wrote the paper. All authors read and approved the manuscript before submission.

## Acknowledgments

We would like to thank the GT QBioS Graduate Program for its support and the Society for the Study of Evolution for granting a Rosemary Grant Advanced Award to Rozenn M. Pineau that helped with pushing this work forward. This work was initiated by a seed grant from AV and JM. The work was further supported by grants from the NIH (Grant No. 5R35GM138030), the NSF Division of Environmental Biology (Grant No. DEB-1845363) to WCR and (Grant No. DEB-1844941) to ZG, and the NSF grant Paleo Perspectives on Climate Change (P2C2) Program (Grant No. 2102997) to JM and AB. The Utah Agricultural Experiment station provided support to KM and this publication is UAES journal paper No. 9835. The support and resources from the Center for High Performance Computing at the University of Utah are gratefully acknowledged.

## Data accessibility

Scripts and detailed steps can be found on the paper repository.

## Supporting information

### Supplementary Methods

We developed a filtering pipeline to isolate the somatic mutation from the germline mutations in our samples.

1. The first step consists in removing common mutations between the dataset of interest and a dataset that could be considered as our ‘normal’. However, we do not have access to the founding genotype. Hence, to account for common mutations and mutations that could have been found in the original seed, we compared the variant files to the (a) set of variants of surrounding clones and (b) a ‘panel of normals’, composed of 100 samples of P. tremuloides from Utah neighboring states (Idaho, Wyoming, Colorado, Nevada) that were collected and sequenced with the large scale dataset in 2008.
2. The vcf file was then filtered for low-quality variants using the following set of filters : minimum coverage per variant set to twice the number of samples, minimum number of sequences with the alternative allele set to 1, filtering out loci fixed for the alternate allele, minimum mapping quality of 30 and maximum number of samples without data set to 80% of the number of samples in the dataset.
3. Next, to calculate point estimates, we calculated the genotype likelihoods and the allele frequency. The expectation of allele frequency for fixation of an allele in a diploid organism is 0.5. However, because the Pando clone is triploid, it reduces our expectation for fixation of a mutation to 0.33. We thus adjusted the allele frequency values in the file to not take into account any variants whose allele frequency *>* 0.7 (set to 0), and reduce to 0.5 any allele frequency comprised between 0.5 and 0.7. We used this modified allele frequency vector to calculate the point estimates. We labeled samples as homozygotes or heterozygotes for every variant by comparing their point estimate value to a threshold value (.95).
4. We filtered out individuals with a mean coverage less than 4x for all variants.
5. To remove variants that may have been present in the founding seed of the organism, we filtered out (a) the SNPs that were found in 80% or more of the samples. We also removed (b) the variants that were only found in one sample, as they could be either rare variants, or sequencing errors.

## Supplementary Figures

**Fig. S1.**
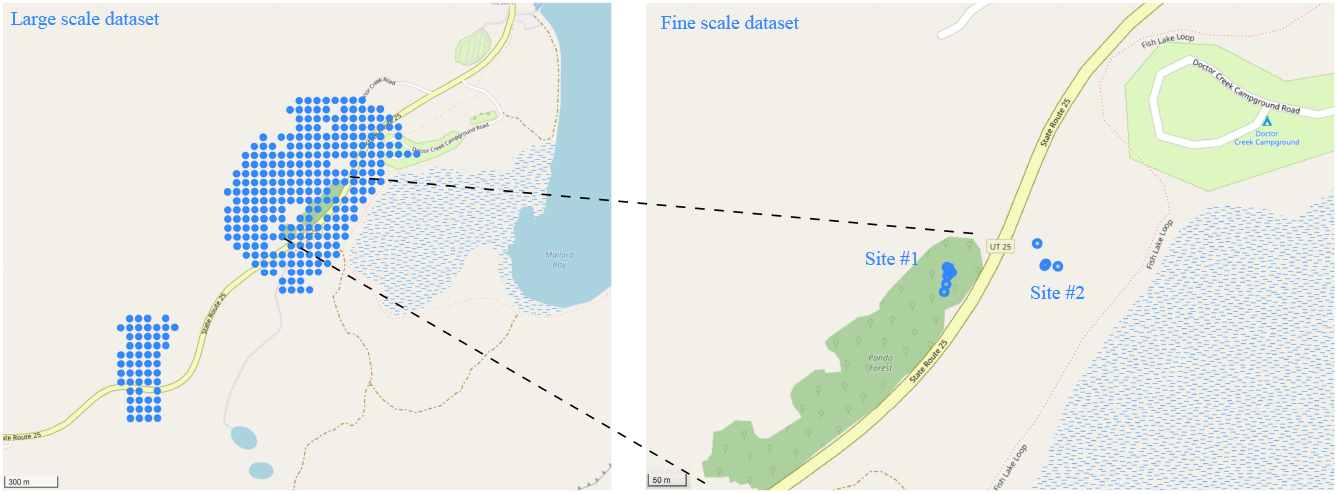
Localities for large-scale (left) and fine-scale (right) sampling. Coordinates are given in Supplementary Table.

**Fig. S2.**
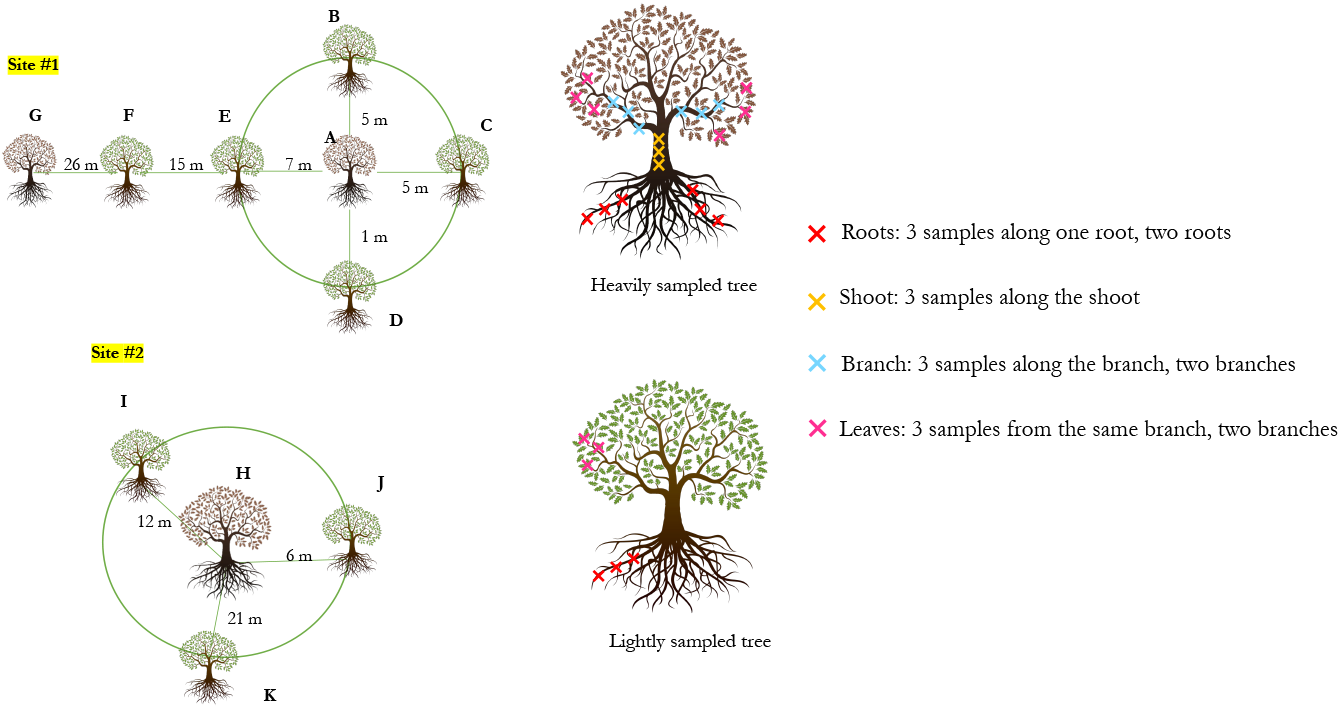
Sampling strategy for the fine-scale dataset. Leaf, bark, branch and root samples from two localities within the Pando stand were collected. In site #1, located in a recently clear cut area, two ramets were heavily sampled (leaf, bark, branch and root samples), and five surrounding ramets were lightly sampled (leaf and root samples). In site #2, one ramet was heavily sampled and three surrounding ramets were lightly sampled.

**Fig. S3.**
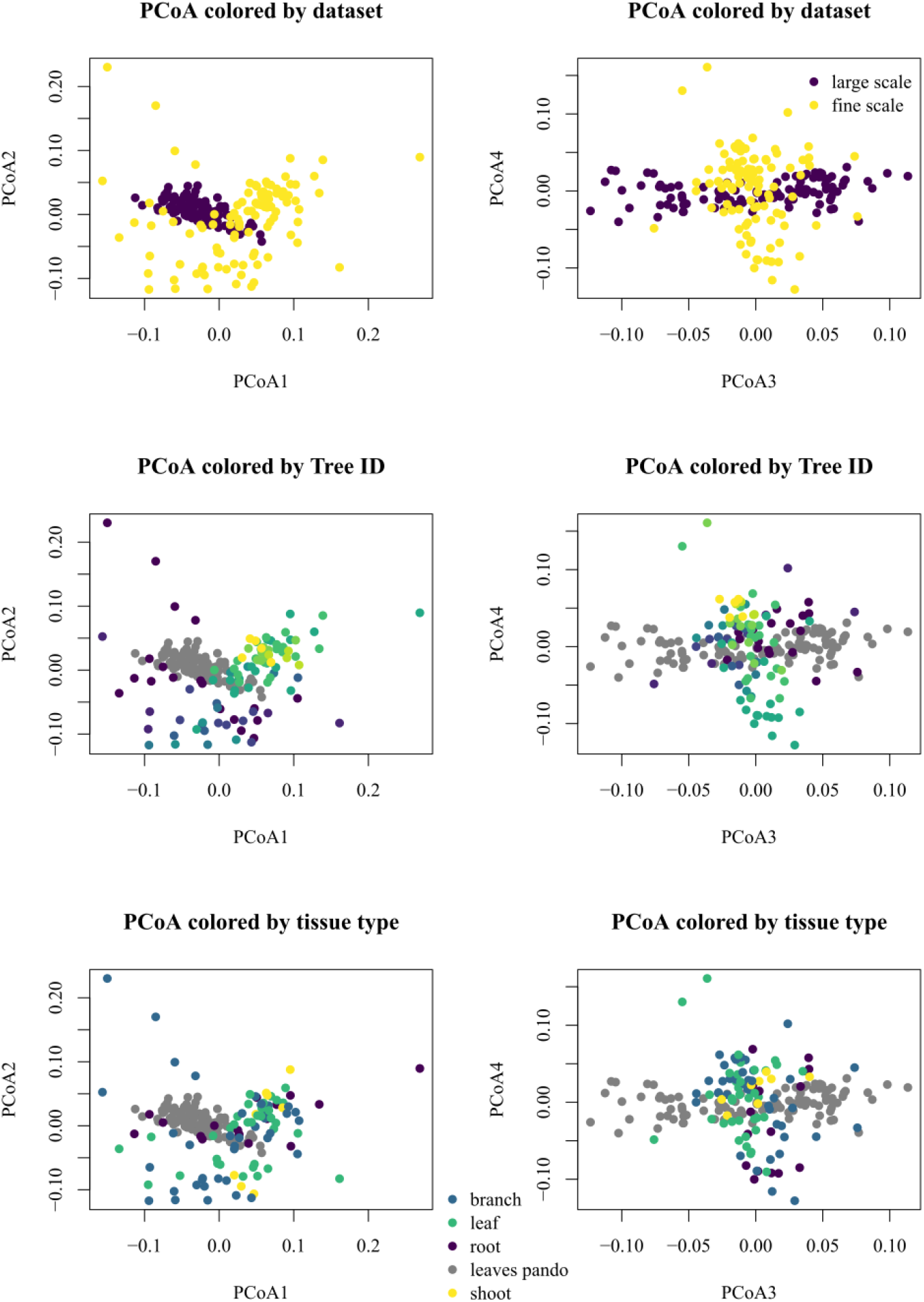
Principal Coordinate Analysis (PCoA) on the large scale and finer scale datasets, colored by dataset, tree ID and tissue type.

**Fig. S4.**
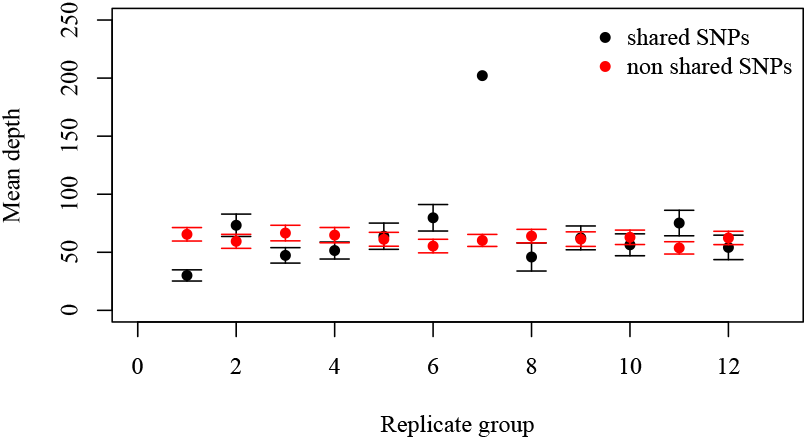
Mean read depth per SNP in the replicate dataset. The mean read depth per SNP for the mutations that were found in more than 2 samples per replicate group was not different from the mean read depth of the mutations that were not found in more than 2 samples per group (two-sided T test, *t* = 0.69, *P* = 0.51). Error bars indicate standard error. Replicate group 7 only had 2 samples left after filtering and was removed from downstream analyses.

**Fig. S5.**
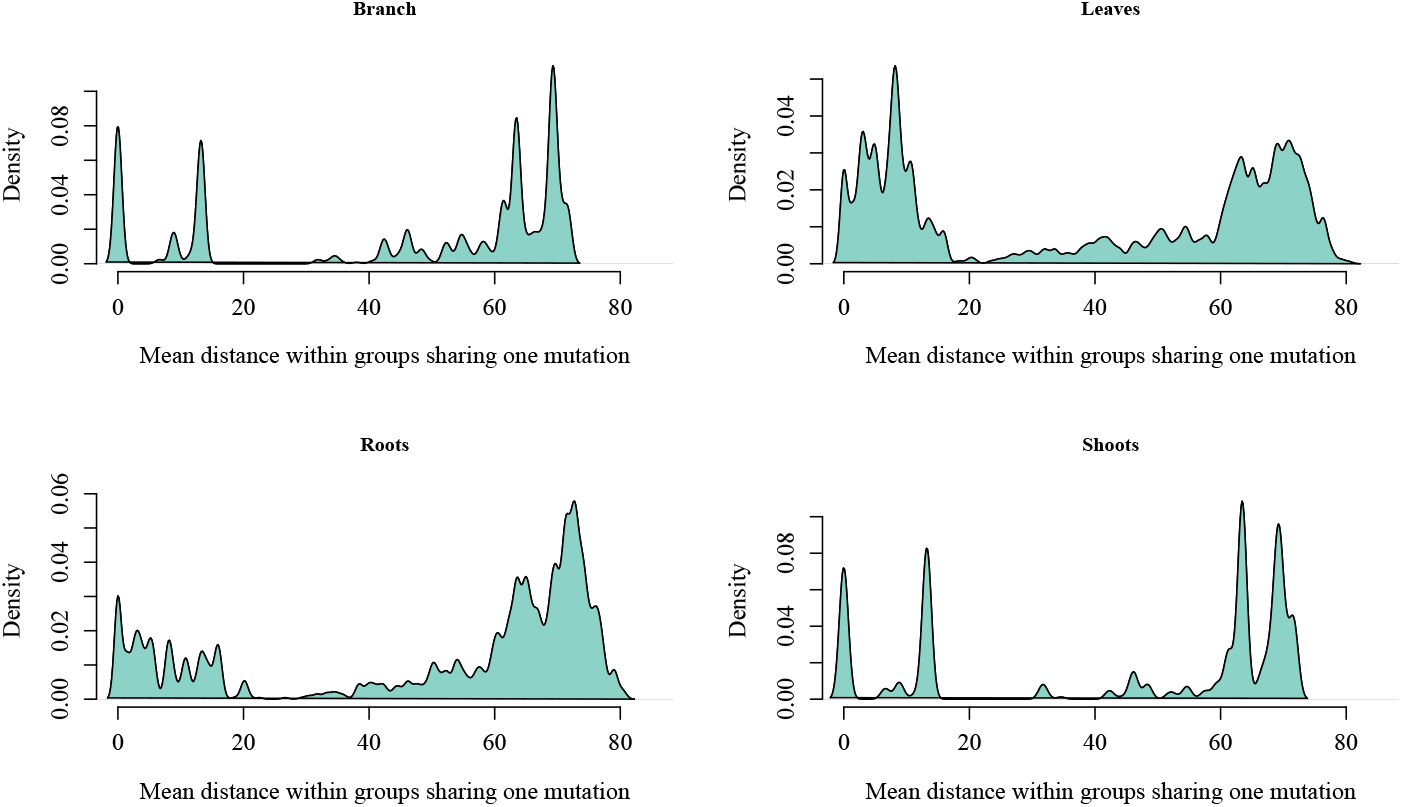
Distributions of mean correlations between the spatial distance between pairs of samples, and the number of mutations they have in common, sorted by tissue type. Correlation values: *r* = −0.06 for branches,*r* = −0.44 for leaves, *r* = −0.11 for roots, *r* = −0.05 for shoots.

**Fig. S6.**
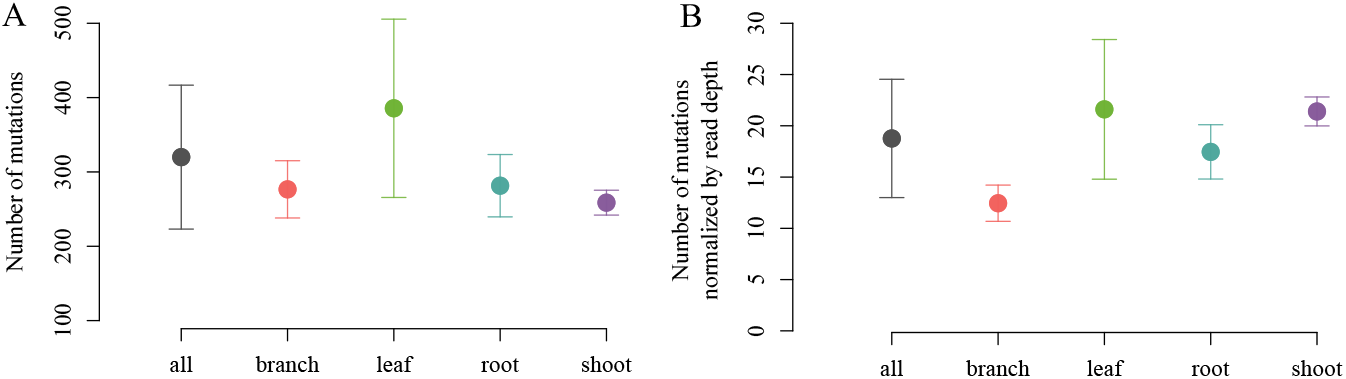
Tissue-specific mutation loads. (A) The number of somatic mutations differs between tissue types (ANOVA, *F*_3,97_ = 14.47, *P* = 7.26*e*^−8^), with the leaves having a significantly higher number of mutations as compared to the roots, branches or shoots (Tukey HSD’s *P <* 0.0003). (B) When normalized by read depth, the leaves still show a significantly higher number of mutations as compared to root and branches, but not shoots (ANOVA, *F*_3,97_ = 16.55, *P* = 9.22*e*^−9^ followed by Tukey HSD with *P <* 0.0001 for roots and branches).

**Fig. S7.**
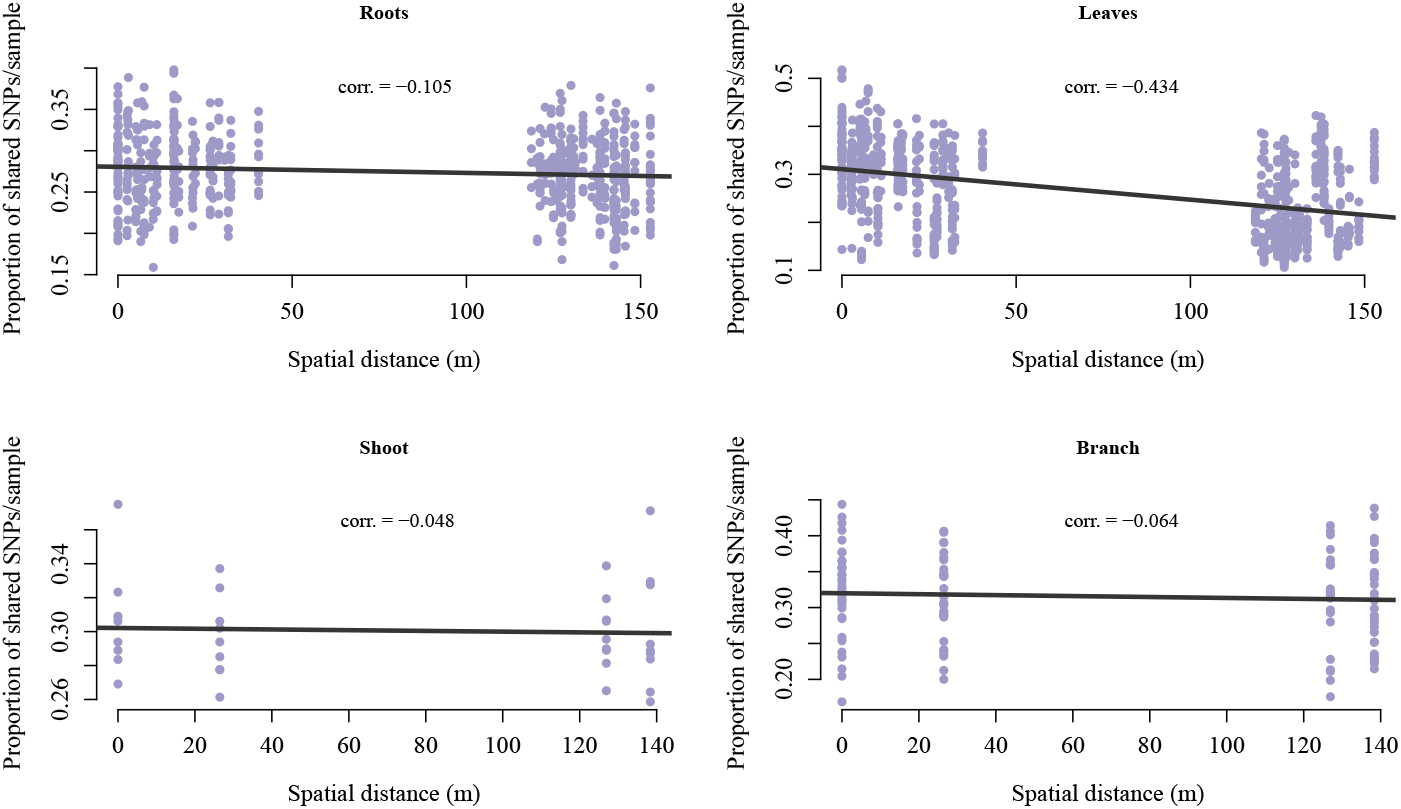
Distributions of mean distance between samples sharing one mutation in the fine-scale dataset, sorted by tissue type. Mean distance for leaves is 39.28 m, mean distance for roots is 51.36 m, mean distance for shoots is 46.12 m, mean distance for branches is 46.69 m.

**Fig. S8.**
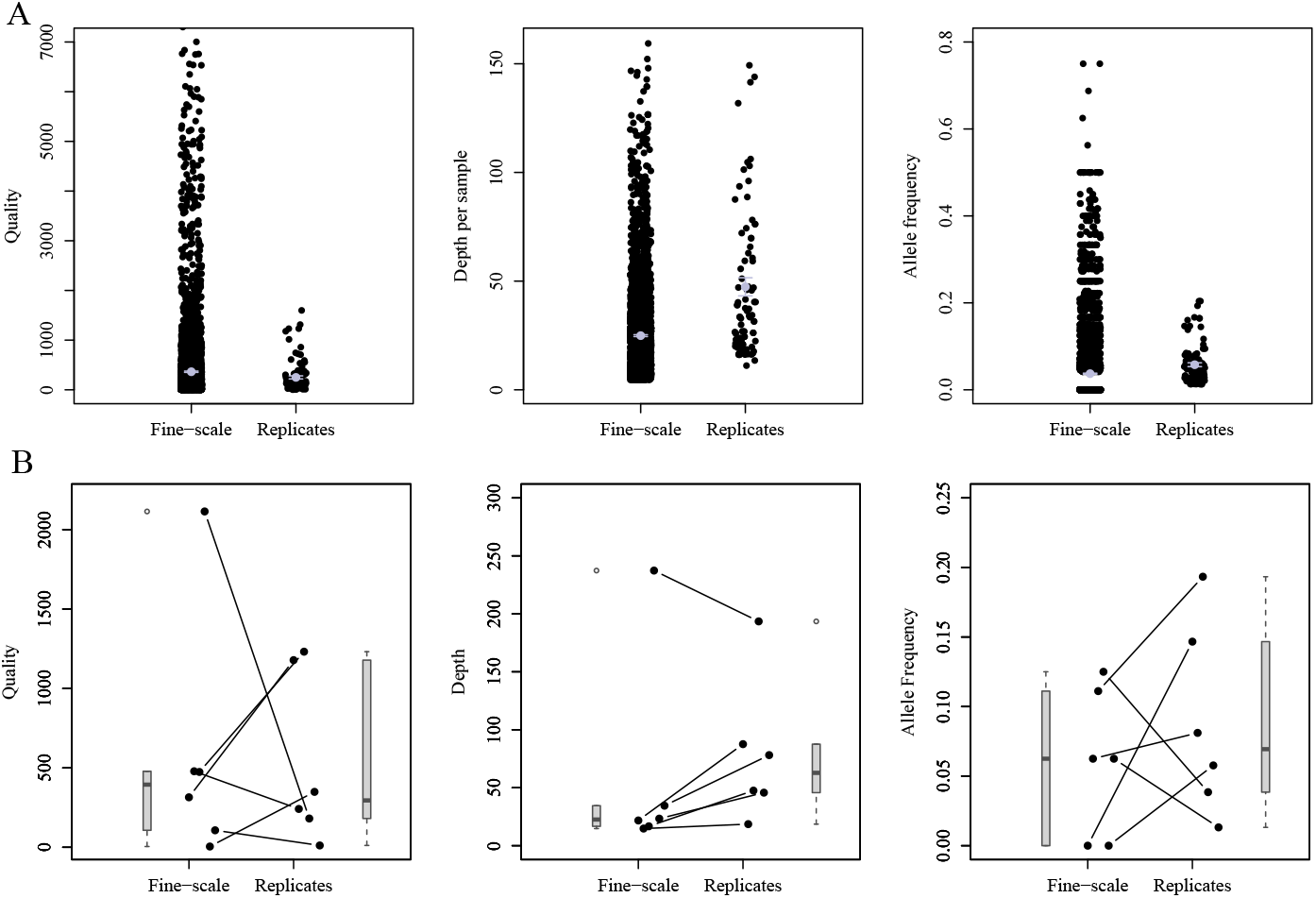
Quality, read depth and allelic frequency comparisons between all sites in the “fine-scale” and the “replicate” datasets (A), and between the set of sites that replicate (B). (A) The mean read depth per sample is overall higher in the “replicates” dataset than in the “fine-scale” dataset (two-sided T test,*t* = −5.37, *P* = 4.93 ×10^−7^). (B) We also observe a higher depth in the set of mutations that replicate between datasets.

**Fig. S9.**
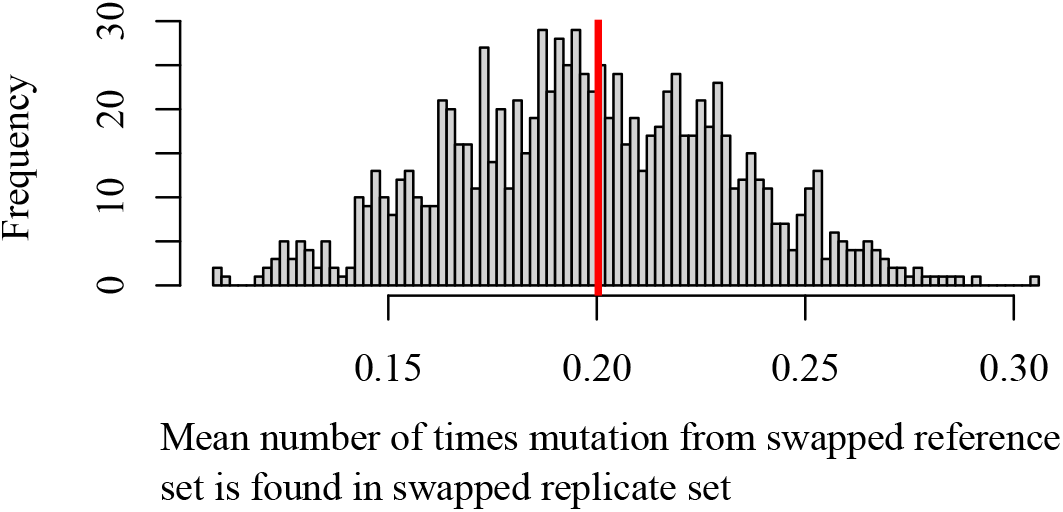
Mean number of times a mutation from the swapped fine-scale sub-dataset is found in swapped replicate set. To test whether the set of mutations from the fine-scale sub-dataset was more likely to replicate than a random set, we isolated one sample at random from the fine-scale sub-dataset (12 samples) and the replicate dataset (8 times these same 12 samples) to create a new reference set. We calculated the mean number of times a mutation identified in a sample from the fine-scale sub-dataset was found in the replicate dataset, and compared to the mean value from the initial replicate and fine-scale datasets (p=0.46).

**Fig. S10.**
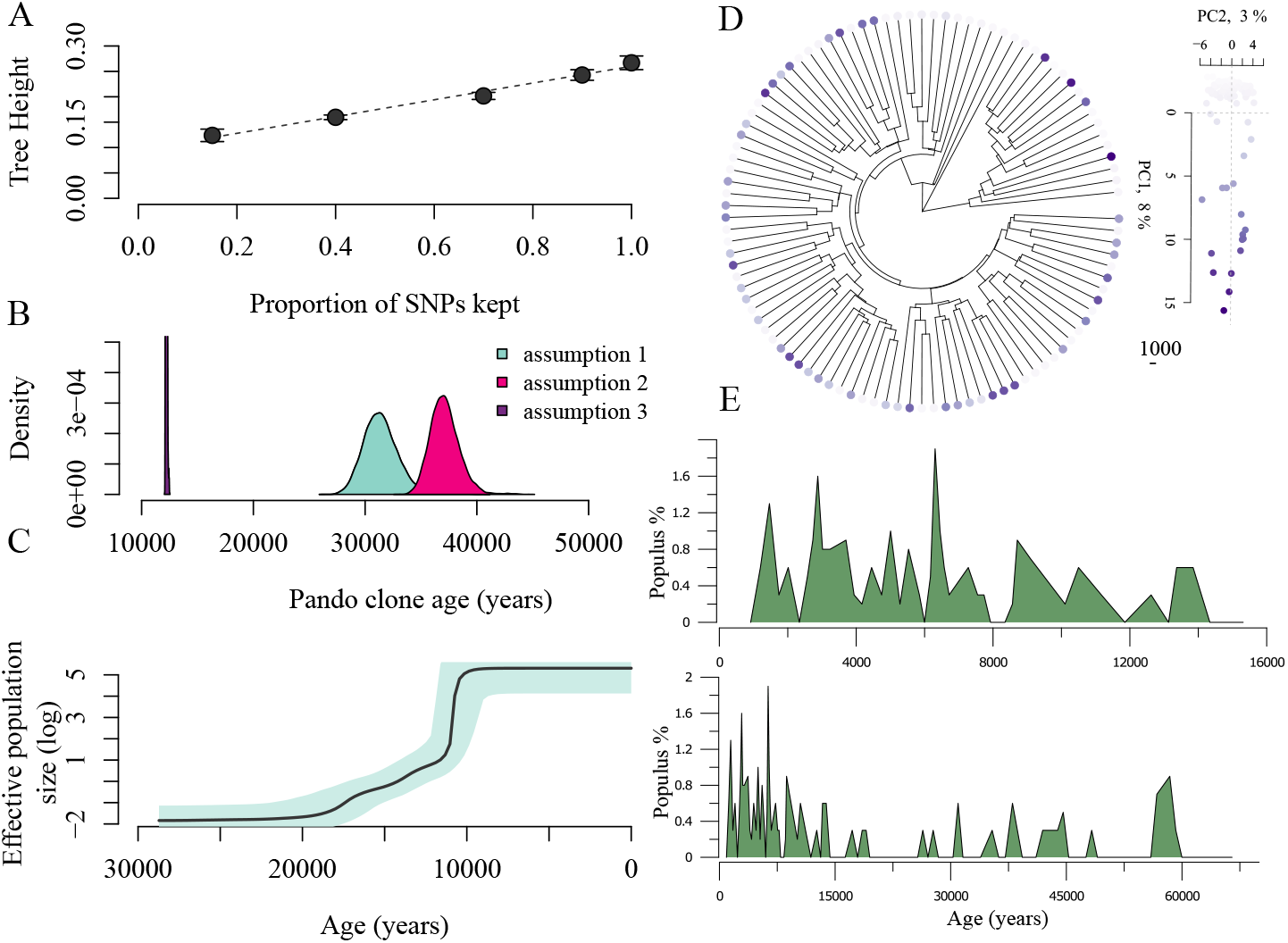
The Pando clone age is between ∼12,000 and 37,000 years old based on the fine-scale dataset. (A) We use the relationship between the proportion of missing mutations from a simulated dataset and the phylogenetic tree height to take into account the somatic mutations that we are missing in the Pando clone fine-scale dataset (linear regression *y* = 0.10 + 0.16*x, P <* 2.2∗ 10^16^, *R*^2^ = 0.95). (B) With this correction, we calculate the Pando clone age based on three different assumptions: (1) if the mutations we detect are all real, the Pando clone would be about 31,421 years old (±sd = 1,584 years); (2) if we are missing 80% of the mutations, then the clone would on average be 37,126 years old (±sd = 1,308 years); (3) finally, if only 20% of the mutations we detect are real somatic mutations, the Pando clone would be 12,215 years old (±sd = 52 years). (C) Despite thousands of years of evolutionary history, the Pando clone shows minimal phylogenetic structure (points colored according to PC1 score).

## Notes

### Competing Interest Statement

The authors have declared no competing interest.

### Summary of Updates

Upload the Supplementary Table cited in the manuscript.

